# Modeling the trajectory of SARS-CoV-2 spike protein evolution in continuous latent space using a neural network and Gaussian process

**DOI:** 10.1101/2021.12.04.471198

**Authors:** Samuel King, Xinyi E. Chen, Sarah W. S. Ng, Kimia Rostin, Tylo Roberts, Samuel V. Hahn, Janella C. Schwab, Parneet Sekhon, Madina Kagieva, Taylor Reilly, Ruo Chen Qi, Paarsa Salman, Ryan J. Hong, Eric J. Ma, Steven J. Hallam

**Affiliations:** Department of Microbiology and Immunology, University of British Columbia, 2350 Health Sciences Mall, Vancouver, BC, V6T 1Z3, Canada; Independent Researcher, Cambridge, MA 02139, USA; Graduate Program in Bioinformatics, University of British Columbia, Genome Sciences Centre, 100-570 West 7th Avenue, Vancouver, British Columbia V5Z 4S6, Canada; Genome Science and Technology Program, University of British Columbia, 2329 West Mall, Vancouver, BC V6T 1Z4, Canada; Life Sciences Institute, University of British Columbia, Vancouver, British Columbia, Canada V6T 1Z3; ECOSCOPE Training Program, University of British Columbia, Vancouver, British Columbia, Canada V6T 1Z3

**Author notes:** Correspondence concerning this paper should be addressed to Steven J. Hallam. Electronic mail may be sent to. Declaration of competing interest: SJH is a co-founder of Koonkie Inc., a bioinformatics consulting company that designs and provides scalable algorithmic and data analytics solutions in the cloud.

**Keywords:** deep learning, Gaussian process, protein evolution, SARS-CoV-2, spike protein, variational autoencoder

## Abstract

Viral vaccines can lose their efficacy as the genomes of targeted viruses rapidly evolve, resulting in new variants that may evade vaccine-induced immunity. This process is apparent in the emergence of new SARS-CoV-2 variants which have the potential to undermine vaccination efforts and cause further outbreaks. Predictive vaccinology points to a future of pandemic preparedness in which vaccines can be developed preemptively based in part on predictive models of viral evolution. Thus, modeling the trajectory of SARS-CoV-2 spike protein evolution could have value for mRNA vaccine development. Traditionally, *in silico* sequence evolution has been modeled discretely, while there has been limited investigation into continuous models. Here we present the Viral Predictor for mRNA Evolution (VPRE), an open-source software tool which learns from mutational patterns in viral proteins and models their most statistically likely evolutionary trajectories. We trained a variational autoencoder with real-time and simulated SARS-CoV-2 genome data from Australia to encode discrete spike protein sequences into continuous numerical variables. To simulate evolution along a phylogenetic path, we trained a Gaussian process model with the numerical variables to project spike protein evolution up to five months in advance. Our predictions mapped primarily to a sequence that differed by a single amino acid from the most reported spike protein in Australia within the prediction timeframe, indicating the utility of deep learning and continuous latent spaces for modeling viral protein evolution. VPRE can be readily adapted to investigate and predict the evolution of viruses other than SARS-CoV-2 in temporal, geographic, and lineage-specific pathways.

## 1. Introduction

The onset of the COVID-19 pandemic, caused by the respiratory disease resulting from infection of the novel coronavirus SARS-CoV-2, fueled a surge in vaccine research and scientific effort across the globe. Despite this effort, it is unclear whether vaccines for SARS-CoV-2 will remain effective over time as the virus evolves into new variants. Currently, the prominent issue is high transmission rate, but over time, the virus may become endemic: which raises the concern of long-term mutational risk combined with waning immunity [1-2]. Moreover, amongst viral genome architectures, single-stranded RNA viruses such as SARS-CoV-2 have the highest rate of mutation per replication, which in part gives them a relatively higher chance of mutating with each infection [3]. This suggests that updated vaccines will need to be carefully designed to compensate for viral evolution between infectious seasons, as is currently the case for influenza vaccinations. Therefore, a key challenge is ensuring vaccine-generated immunity remains effective even after viruses undergo multiple mutation cycles [4]. To establish long-lasting immunity in this context, it is crucial to have predictive platforms in place to allow the preemptive study of variants, inform vaccine development, and increase pandemic preparedness.

Currently, mRNA vaccines for SARS-CoV-2 specifically target the spike protein [5]. The spike protein is the surface glycoprotein that enables SARS-CoV-2 entry into human cells through binding of the angiotensin-converting enzyme 2 (ACE2) receptor [6-8]. The importance of the spike protein for host cell entry makes it a logical target candidate for vaccines, and therefore modeling spike protein evolution could inform vaccine development. Attempts to model viral evolution *in silico* have typically been carried out using computational models that aim to simulate general mechanisms of natural evolution, such as genetic drift [9]. For example, the arrival of new influenza strains each year has encouraged the development of various predictive platforms [e.g., 10-11], which are useful in the development of annual flu vaccines [12].

Newer models for predictive evolution, particularly those involving machine learning, tend to extrapolate patterns from recorded evolutionary events, rather than impose theorized patterns of natural processes [13]. Deep learning is a subfield of machine learning that uses neural networks to learn from large amounts of data [14]. Deep learning is emerging as a promising method for modeling biological processes and has already been used to model SARS-CoV-2 evolution in a discrete context [15-17] and through natural language models [18]. Of the different types of neural networks, variational autoencoders (VAEs) are viable candidates for amino acid sequence processing due to their ability to reduce multidimensional data down to numerical values that still capture biologically relevant features [19]. VAEs are an unsupervised deep learning approach, meaning that their patterns and features are extracted from unlabeled training data. In the case of SARS-CoV-2 spike protein evolution, a VAE can compress amino acid sequences into continuous latent space, which is a continuum of numerical values where similar values are located closely together. As a type of generative model, VAEs are often used for applications such as image and text generation [20-21], but they have also been used to capture biological information such as sequence mutations and novel cancer biomarkers [22-23]. Through numerical encoding, VAEs can be used to reduce protein sequences into lower dimensions, where they can be further analyzed for evolutionary patterns that are otherwise difficult to detect [24].

When paired with probabilistic models, such as Gaussian processes (GPs), the continuous coordinates encoded by a VAE can be fitted into evolutionary trajectories and projected forward in time. GPs are a Bayesian learning technique that construct probability models of previously observed data, which they can make inferences from [25]. GPs have been used to model various protein properties (e.g., ligand-binding affinity or enzyme activity) with high quantitative accuracy [26]. Regression modeling and time-series forecasting are common applications of GPs. For time-series regression, a GP can fit functions to a given set of data and timepoints and generate regression functions with associated probability distributions that allow for modeling of temporal trends [27]. GPs present a significant advantage over discrete models because their predictions are not restricted to discrete data, and they can predict protein properties across a great diversity of sequences [26]. The ability of GPs to quantify uncertainty helps to determine the validity of the outputs, and utilizing them to model temporal trends also makes fewer assumptions on the shape of data distribution [26, 28], thus providing a more reliable model. Previous studies have shown the utility of GPs and/or latent spaces to study phylogenetic relationships, model protein stability, design proteins, and in inferring chemical species involved in biochemical interaction networks, among other uses [22, 29-32]. However, the use of GPs and latent spaces for predicting unseen evolution on labelled timelines has not been employed. As a whole, when combined with the generative learning of VAEs, the predictive feature of GPs makes for a synergistic framework for modeling viral protein evolution.

Here, we describe the Viral Predictor for mRNA Evolution (VPRE), a novel approach to protein sequence prediction which models spike protein evolution as a continuum of numerical coordinates, rather than as a discrete timeline of amino acid sequences (Fig. 1). VPRE encodes SARS-CoV-2 spike protein sequences with a VAE and uses a GP to project the most statistically likely chronological trajectories of spike protein evolution. As a proof-of-concept, we made predictions one, two, and five months into the future using validation data from Australia. Our model predicted six putative spike proteins that closely resemble the composition of spike proteins that appeared in real time, differing by only 0 to 3 amino acids depending on the sequence. The nearest neighbor (or the most common sequence differing by 1 amino acid) of the most frequent prediction was also the most frequent spike protein in real sequences from the same time period. These data suggest that modeling protein evolution in continuous latent space holds promise for predictive vaccinology.

**Figure 1.**
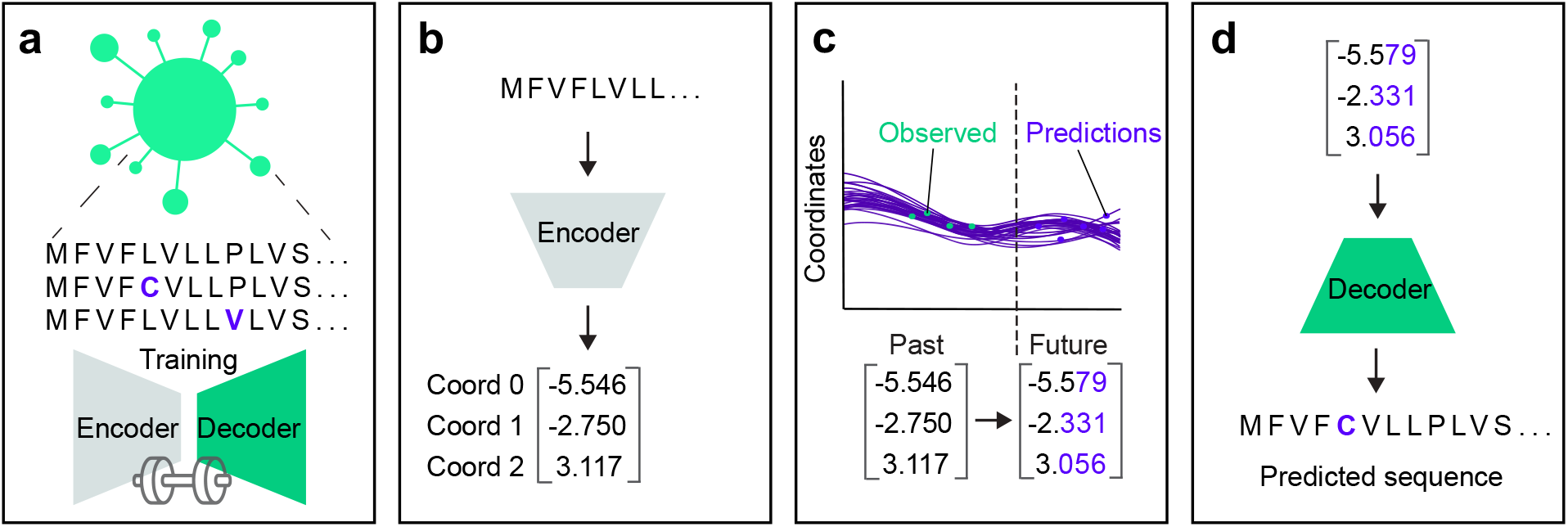
Overview of the VPRE workflow. (a) Step 1: Profile and diversity analysis of over 9000 SARS-CoV-2 spike protein sequences. Based on the diversity profile, the VAE trains on a set of 20,000 simulated spike sequences. (b) Step 2: each sequence in the data set is encoded into three continuous numerical variables, or coordinates, that represent spike proteins from the training data. (c) Step 3: The GP regression finds the best statistical fit of the coordinates over time, and projects probable future coordinates. (d) Step 4: Projected coordinates are decoded into putative sequences resulting from SARS-CoV-2 evolution.

## 2. Materials and methods

### 2.1. Data and code availability

This project was carried out by the 2020 University of British Columbia International Genetically Engineered Machine (iGEM) team, during the period of May 2020 to June 2021. Code associated with this work can be found in our GitHub repository: https://github.com/UBC-iGEM/VPRE.

### 2.2. Data acquisition and construction

The VPRE training dataset consisted of 9534 SARS-CoV-2 spike protein sequences from around the world at varying time points throughout the pandemic (Suppl. File 1). These were downloaded along with their corresponding metadata from NCBI Genbank on August 13, 2020. The five-month VPRE validation dataset from Australia consisted of 8488 spike protein sequences, which were downloaded on January 15, 2021. Incomplete sequences containing gaps (dashes and asterisks) and sequences with ambiguous amino acids (null, B, Z, X) were removed to ensure high quality data.

A training set of 20,000 semi-random mutated spike protein sequences was generated algorithmically by ensuring equal representation in the dataset of any amino acid substitution that occurred at least one time in the spike protein dataset. The algorithm started with the consensus sequence of our analyzed sequences, and went through each position stepwise, presenting with equal likelihood any point mutation observed in the data. This was repeated until 20,000 unique sequences had been created, with the effect of amplifying the presence of infrequent point mutations so that they could be more accurately decoded from predictions made by the GP. No change was made to the analyzed dataset, which remained as sequenced, and only underwent a multiple alignment prior to processing by the GP. The training dataset maintained all the same conserved regions as the real-world data.

### 2.3. Encoding and decoding amino acid sequences through a VAE

As a preprocessing step, training set sequences were aligned by progressive alignment via the Multialign function in the MATLAB bioinformatic toolbox [33]. The aligned sequences were padded with asterisk (*) characters to maximal length and one-hot encoded in order to yield binary matrix representations of the SARS-CoV-2 spike protein sequences that could be inputted into our deep learning model.

The VAE consisted of an encoder and decoder network, where the encoder compressed the sequence data (one-hot encoded amino acid sequences) to its latent embedding and the decoder decompressed the sequence data from its latent embedding. The VAE was implemented in Keras (version 2.4.0) [34] using a TensorFlow backend (version 1.4.0) [35]. In the encoder, the number of latent dimensions was set to three to allow for easier visualization, and thus each sequence was compressed to three numerical coordinates. The latent space distribution was defined with a latent mean and logarithmic variance. Both were calculated with Keras Dense with one-hot encoded training and with an input dimension of three. A standard sampling layer and a Dense layer were created in the encoder. The sampling layer randomly sampled data from latent space following a normal distribution with a mean of zero and a standard deviation of one. The Dense layer mapped the sampled data points to the latent distribution. The decoder was constructed using the encoded data as input and the last layer of the autoencoder as output.

The VAE model was compiled with an Adam optimizer and custom-built loss function. The loss function was the sum of a reconstruction term and a regularization term (expressed as the Kullback-Leibler divergence between the distribution returned by the encoder and a standard normal distribution):

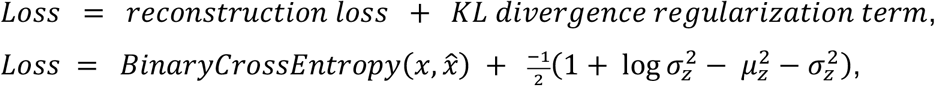

where *x*represents the input data, 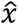 represents the reconstructed data, 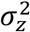 represents the variance of the latent distribution, and *μ*_*z*_represents the mean of the latent distribution. The reconstruction loss served as a measure of the efficacy of the encoder-decoder, as it represented the difference between the reconstructed (decoded) sequences and the input sequences (calculated using binary cross-entropy). Over training, the reconstruction loss was ultimately minimized. The regularization term helped in learning well-formed latent spaces and reducing overfitting during the training process [36]. An early stopping function was also applied with a patience parameter of two in order to stop training once the validation loss metric had stopped improving for two consecutive epochs, thus avoiding overfitting.

### 2.4. Modeling the trajectory of spike protein evolution with GP regressions

A GP was used to model temporal trajectories of each coordinate of Australian sequences encoded by the VAE. After removing duplicated sequences from each day to simplify the model, 185 sequences were obtained (Suppl. File 3). Data from sequences that were collected up to May 31, 2020 (n = 104) were used as a training dataset for the GP, and the rest from June 1 to July 31 (n = 81) were used for validation.

The PyMC3 package (version 3.11) [37] was used to construct the GP model. To model a temporal axis in the GP, an array was constructed to represent the number of days since the first sequence collection in Australia. The other axis in the GP consisted of the coordinate values from the VAE.

The GP was defined as *Y* ∼ *GP*(*K*(*x,x*^′)^,μ(*x*)), adapted from Eric Ma’s Flu Forecaster (https://github.com/ericmjl/flu-sequence-predictor/blob/master/flu-forecaster.ipynb) with a GP latent variable implementation sample [37].

First, the covariance function was defined as an exponentiated quadratic function. The exponentiated quadratic kernel is a popular kernel used in GP modeling, thus it was chosen as a starting point for modeling the data. Because the VAE coordinates were modeled individually, an input dimension of 1 was used for the exponentiated quadratic kernel. The GP model was computed as follows:

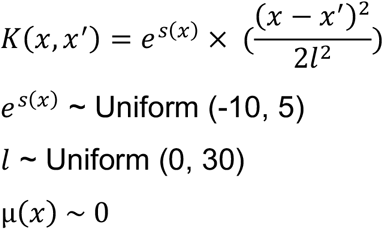

The Uniform function in PyMC3 was used to construct the exponent and Theano was used to construct the exponentiation, followed by a deterministic transformation by using the Deterministic function from PyMC3 [37-38].

Second, a Student’s T log-likelihood distribution was defined to model uncertainties in the covariance function and the input data, adapted from a PyMC3 tutorial (https://docs.pymc.io/notebooks/GP-Latent.html):

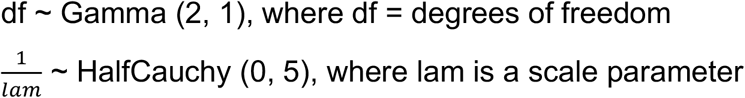

Lastly, the covariance and the mean function were assembled in a Latent GP model. The Exponentiated Quadratic covariance function and the time array were defined for a Latent GP. A Uniform log-likelihood distribution was applied to describe length-scale, as well as a HalfCauchy distribution and a Gamma distribution to define the uncertainty in the covariance function and to model the noise. The VAE coordinates were inputted as observed prior.

### 2.5. Extrapolating the trajectory of GP models

To extrapolate the trajectories the GP predicted, a new time array was set from 0 to 120 + x, where 120 was the number of training days and x represented the number of days into the future to predict. The variable x was chosen as x = 30, 60, and 150 to predict one, two, and five months into the future. The new time array was applied on the GP and a conditional distribution of the predicted functions was obtained with the new input time values using the *conditional* function. 1000 samples were drawn from the GP posterior for each of the three VAE coordinates and merged into 1000 triplets to represent the predicted numerical representations of spike proteins. The triplets were decoded by the decoder of the VAE to obtain predicted spike proteins. The likelihood of each sequence existing in the predicted timeframe was estimated based on the fraction of the 1000 GP predictions that translated exactly to the sequence. Additional packages used in the pipeline include numpy (version 1.19.5) [39], SciPy (version 1.4.1) [40], and pandas (version 1.1.5) [41].

## 3. Results

### 3.1. Amplifying spike protein dataset variation and training the VAE

We used a VAE to compress spike protein sequences and create a biologically relevant latent space. Our dataset included 9534 spike protein sequences collected between January 25 and June 30, 2020. An initial obstacle when training the VAE was that the limited diversity in our relatively small dataset made it difficult for the neural network to identify patterns due to class imbalance (Fig. 2a, b). Across all sequences, the average mutation frequency per amino acid variant was 1.5 × 10^−4^ (Fig. 2d). As expected, we found that the spike protein is quite conserved, especially in its receptor binding domain, where mean variant frequency per amino acid was 6 × 10^−5^ (Fig. 2c). To improve the VAE training, we simulated 20,000 spike proteins with an amplified but equal chance of mutation at any of the mutation sites seen in the original protein sequences, while the conserved regions were maintained (Fig. 2e). The majority of variant frequencies at each position consequently rose to approximately 0.3 or 0.5, given that there were many amino acid positions with two or three variants. This approach increased the VAE’s ability to encode and decode rare variants in the dataset, which ensured that low-frequency variants were represented in the GP input and could be decoded accurately if predicted.

**Figure 2.**
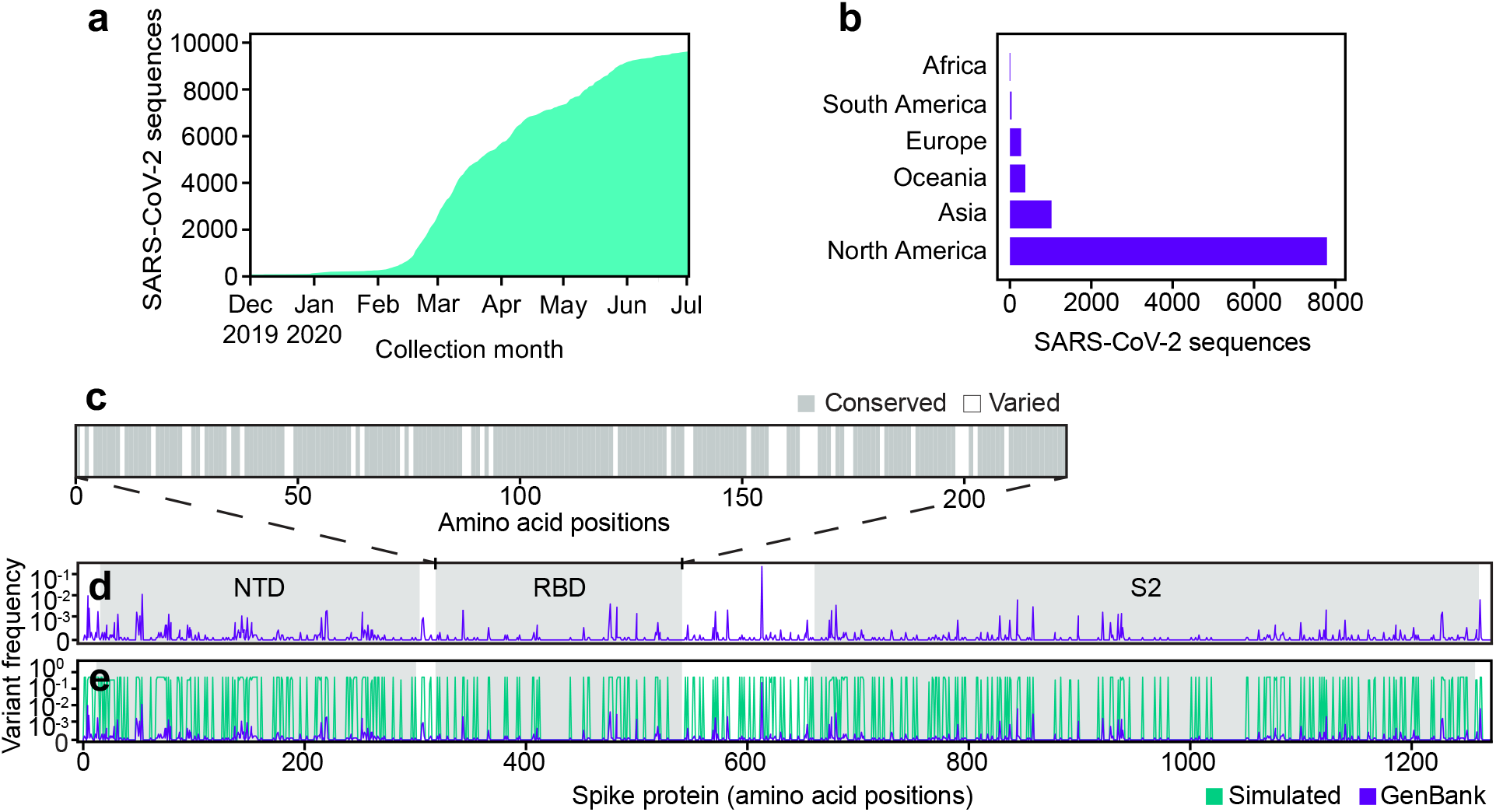
The collected and simulated datasets for neural network training and encoding. (a) Cumulative distribution of sequences over time from the dataset downloaded from NCBI GenBank on August 16th, 2020 (n = 9534). (b) Continental distribution of sequences from the GenBank dataset. (c) Distribution of amino acid variations in the spike receptor binding domain (white = at least one variant observed from the GenBank dataset; purple = no variant detected in the GenBank dataset). NTD, N-terminal domain; RBD, receptor binding domain; S2, spike protein S2 domain. (d) Variant frequency at each amino acid position on spike proteins observed from the NCBI dataset. (e) Comparison of variant frequency at each amino acid position on spike proteins in the GenBank dataset (purple) and the simulated dataset (green).

We trained the VAE for 41 epochs, determined by an early stopping function, where a single epoch included one round of encoding and decoding. This was followed by the calculation of a difference score between the input and output sequences, which represented the loss or error of the model (Fig. 3a,c). As a simple means to verify whether or not the sequence encodings outputted by the VAE accurately captured differences in the spike protein sequences, the Euclidean distances between the numerical latent coordinates were compared to the Levenshtein distances between the amino acid sequences (Fig. 3d). A Euclidean distance is the length of a line segment between two data points in geometric space, while a Levenshtein distance is a metric for measuring the number of differences between two strings [42]. The two variables were strongly correlated (r = 0.79), suggesting that sequences became increasingly different as their distance in latent space increased, and that variation within amino acid sequences was well-captured in the VAE-encoded numerical coordinates.

**Figure 3.**
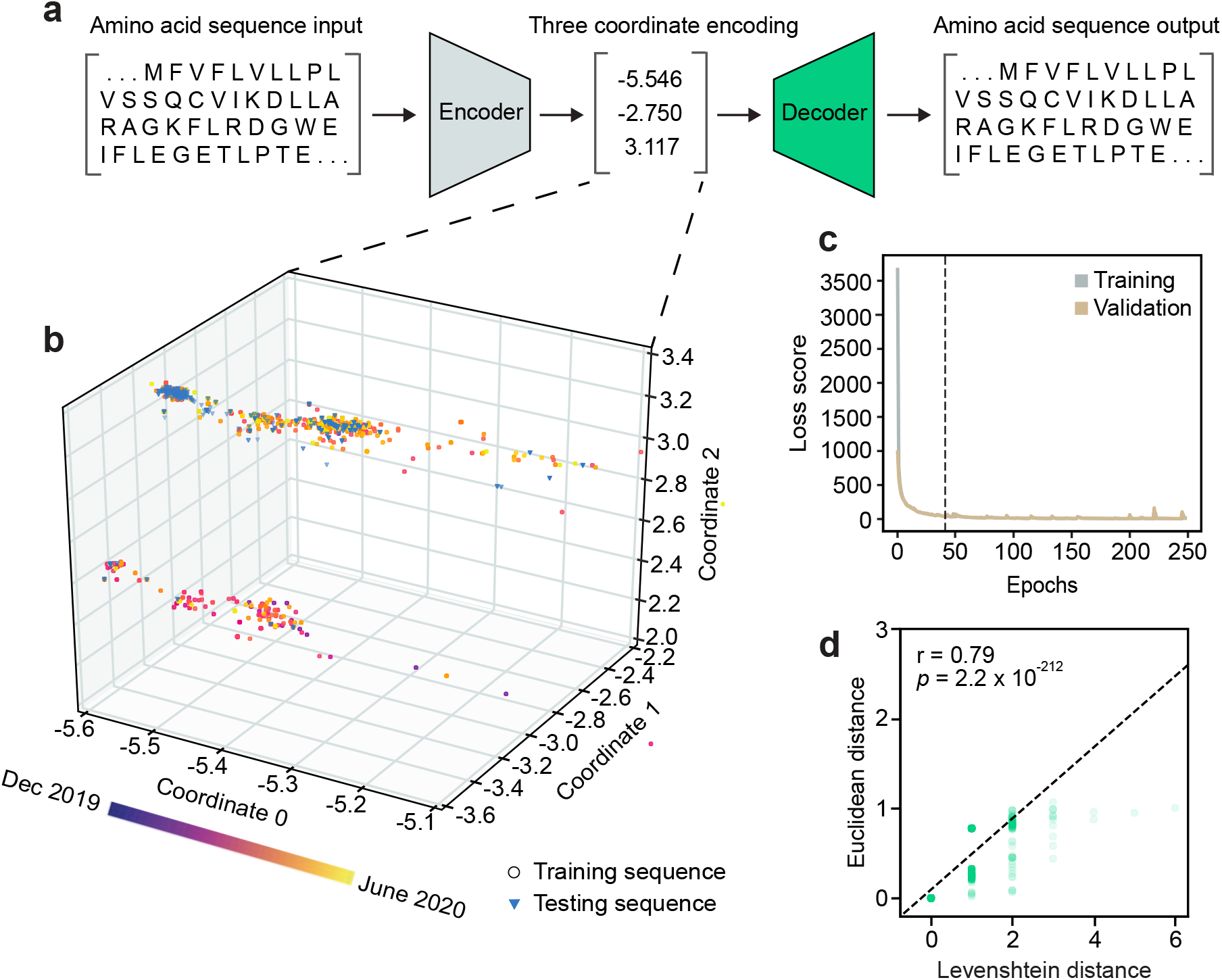
The architecture and unsupervised learning of the VAE. (a) Illustration of the VAE architecture. Three latent dimensions, or coordinates, were set for VAE-translated variables. (b) Overview of latent representations of the viral spike protein sequences. Sequences collected prior to May 30, 2020, are grouped as a testing dataset and are represented by circles (n = 7620). Sequences collected after May 30, 2020, are grouped into a validation dataset and are represented by triangles (n = 1914). (c) VAE training loss curves. The dashed line indicates the number of epochs used in training the final model. (c) Correlation of Levenshtein distances of each sequence pair in the NCBI dataset and Euclidean distances of corresponding latent representations from the VAE. The black dotted line is the fitted line. The significance threshold was adjusted by Bonferroni correction. n = 7620.

### 3.2. Encoding spike proteins into continuous latent space using the trained VAE

After training the VAE on 20,000 simulated sequences, we encoded 7620 spike protein sequences collected before May 30, 2020, into three latent dimensions, or numerical coordinates (Fig. 3a,b). The continuous latent space representation created by our sequence encodings separated into two major populations: one proximal to the sequences collected near December 2019, and the other proximal to those from May 2020 (Fig. 3b). We used the remaining 1914 sequences from the dataset collected after May 30, 2020, to validate the latent space generated by the first sequence encodings. As expected, the validation data appeared in the latent space within the population of sequences proximal to May 2020. This suggests that the VAE could learn a latent variable model and the parameters of the probability distribution modeling the input data.

### 3.3. Modeling and predicting evolutionary trajectories using GPs

To forecast novel predictions, we inputted each coordinate of the encoded sequences from the VAE into individual GPs. Each GP performed a regression analysis on the input coordinates from the VAE to find the best fitting functions to the data points in chronological order. After this training period, the functions were projected into the future to predict the sequences of the most statistically likely spike proteins that might evolve based on previous evolutionary patterns. Because GP predictions are continuous coordinates and amino acid sequences are discrete, multiple coordinate triplets can represent the same amino acid sequence. As a result, a frequency index can be calculated for each predicted sequence, which we used to estimate their likelihood.

As a proof-of-concept, we tested the ability of GPs to predict spike protein evolution by training on sequences from Australia collected prior to May 30, 2020 (n = 104), and projecting the trajectories of 1000 sequences one, two, and five months into the future (Fig. 4). We chose Australia with the assumption that it would allow us to simulate an isolated and simplified phylogenetic pathway for SARS-CoV-2 spike proteins, under the hypothesis that Australia as an island continent was more isolated and therefore, less subject to external factors contributing to variant emergence. Moreover, at the onset of this work Australia had the most sufficient spike protein data available for our analysis when compared to other island nations. Within the training period, the functions of all three coordinates tightly fitted with the training sequences. In the first two months of predictions, the range of coordinate values expanded, but then stabilized throughout the five-month prediction period. This can also be seen in the frequency distributions for each coordinate and their respective prediction periods, where the predicted values for each coordinate generally followed a normal distribution, and months two and five appeared to have similar value distributions. Clear clustering of the training data points is seen in Coordinates 1 and 2, suggesting the presence of two dominant spike protein sequences within the Australian dataset.

**Figure 4.**
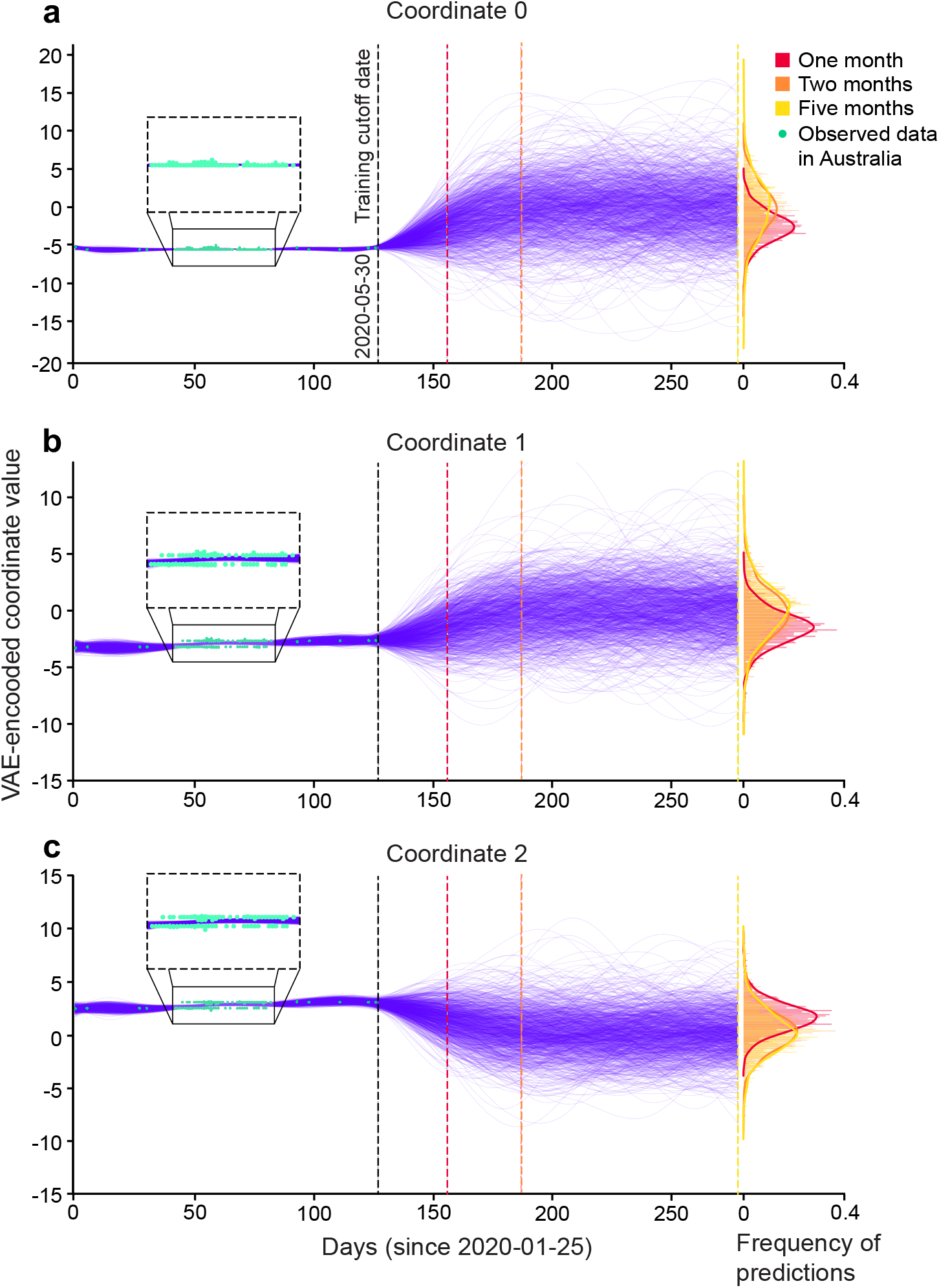
Spike protein sequences from Australia projected up to five months into the future in the GP regression. Trajectories of latent representations of sequences analyzed by the GP (purple lines; n = 1000) with the training coordinates overlaid on the training period (green dots; n = 104). After training the GP on sequences up until May 30, 2020 (n = 104), predictions were made one month (blue dotted line), two months (red dotted line), and five months (yellow dotted line) into the future. The corresponding frequency distributions of each prediction period are shown on the right.

The VAE decoded the 1000 coordinate triplets predicted by the GP at the end of five months into 17 amino acid sequences (Fig. 5a). Upon a BLAST analysis, we found that the top two predictions were existing spike proteins and the other 15 were novel sequences. To investigate whether the model produced unseen amino acid variants, we compared the 17 predicted sequences against all training sequences (Fig. 5b). We found five novel amino acid substitutions and four deletions that were only seen in the predicted sequences and not in the training data. Most of the unseen amino acid variants occurred at variable regions in the training set. Interestingly, there were three predicted amino acid variants at conserved regions within the training data. These results indicate that the model was able to produce sequences other than those that it was trained on.

**Figure 5.**
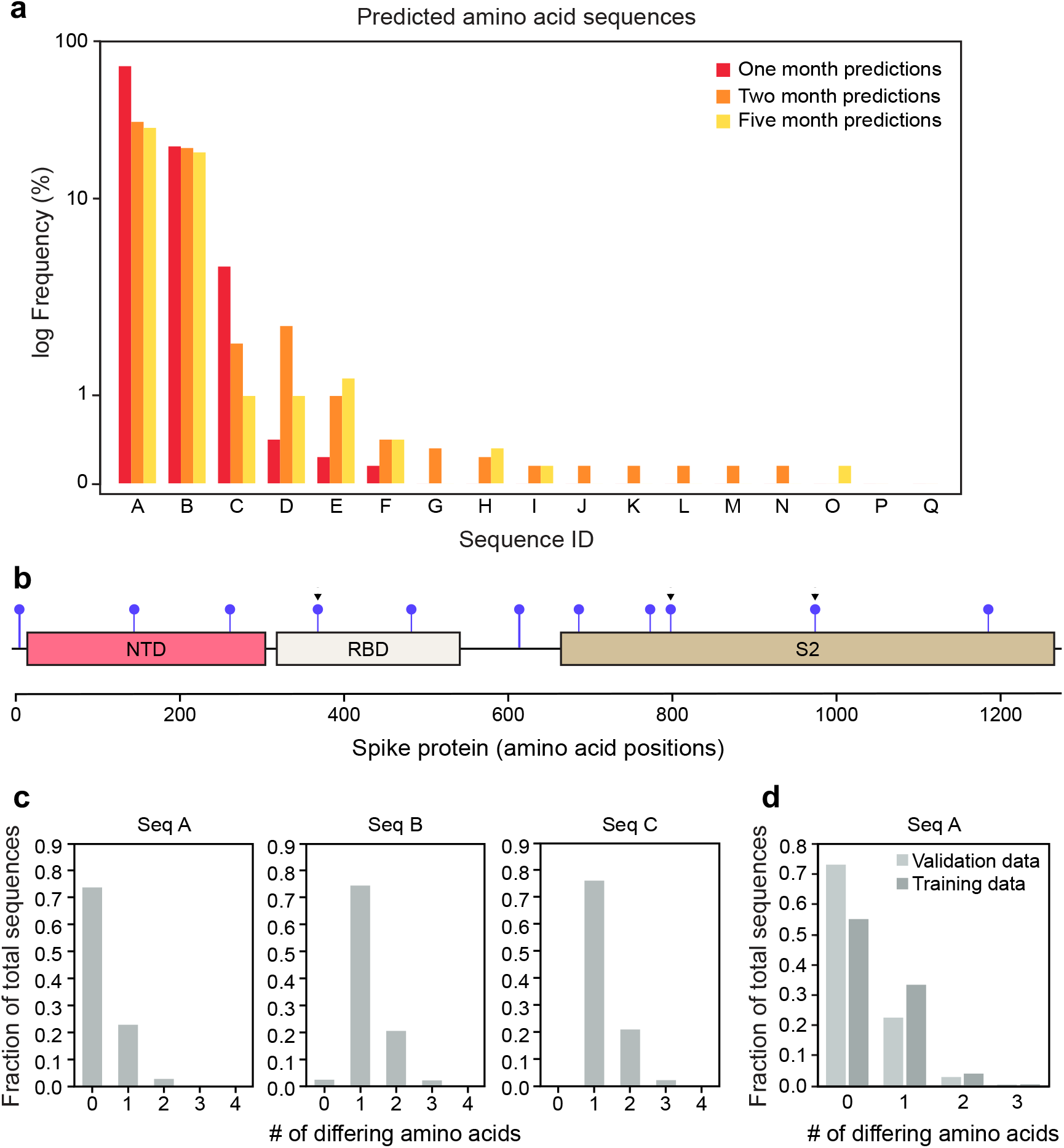
Spike protein variant predictions from the GP. (a) Predicted amino acid sequences decoded from three GP regressions performed on coordinates 0, 1, and 2. The frequency of predictions is ordered from highest to lowest from A to Q. (b) Unique variant positions produced by the VAE that were unseen in the training dataset, indicated by purple lollipops. Arrowheads indicate conserved positions in the training dataset. Novel variants include deletions. (c) Number of amino acid differences between the top three predictions and validation sequences collected in Australia up to one month after May 30, 2020 (n = 81). (d) Number of amino acid differences between the most frequent prediction (Seq A) and the GP training sequences (n = 104) and validation sequences (n = 81).

Across the 17 predicted sequences, six persisted across all five months of predictions. We performed further analysis on the top three predictions due to their high frequency out of the 1000 GP predictions (Fig. 5c; Suppl. Fig. 2). We did not consider the remaining 14 sequences as strong prediction candidates. However, for these low-frequency predictions, the top matches from BLAST analysis showed that they still had over 99% identity and query coverage to SARS-CoV-2 spike proteins.

As a control, we compared the top three predictions to the sequences collected in Australia one month after our training data cut-off date of May 30, 2020 (n = 81, Fig. 5c). The top prediction comprised over 70% of the validation dataset, while the second- and third-most probable predictions were identical to less than 25% and 5% of the validation sequences, respectively. When comparing the top prediction to our GP training sequences (n = 104), around 55% were identical to our top prediction (Fig. 5d). This suggests that our GP worked as expected even when trained on a small dataset, given that it predicted the most dominant sequence it was trained on, which was also the most dominant sequence present in Australia during the prediction period. The top two predictions being identical to existing spike proteins in Australia also suggests that the VAE could reproduce accurate spike protein sequences.

We retrieved 8407 additional Australian spike protein sequences collected between the training cut-off date of May 30, 2020, and November 30, 2020 (the five-month prediction period) and compared them against our three most frequent predictions (Fig. 6a). Over 85% of the newly retrieved sequences differed from our most frequent prediction by only one amino acid (N477S), and over 90% of these nearest neighbors were identical to each other. The most common nearest neighbors of the second- and third-most frequent predictions differed by only two amino acids from the predictions. When tracing the frequency patterns of the two sequences in our Australian dataset between January and November 2020, we found that the top prediction was the most prevalent spike protein up until May (Fig. 6b). After May, the top prediction’s nearest neighbor became the most prevalent spike protein in Australia, emerging in April, reaching a frequency of around 90% by July and outcompeting our predicted sequence. Taken together, the GP’s top prediction was off by a single amino acid when extrapolating the most dominant spike protein five months into the future. This suggests that our model could make meaningful predictions even when trained on only 104 amino acid sequences.

**Figure 6.**
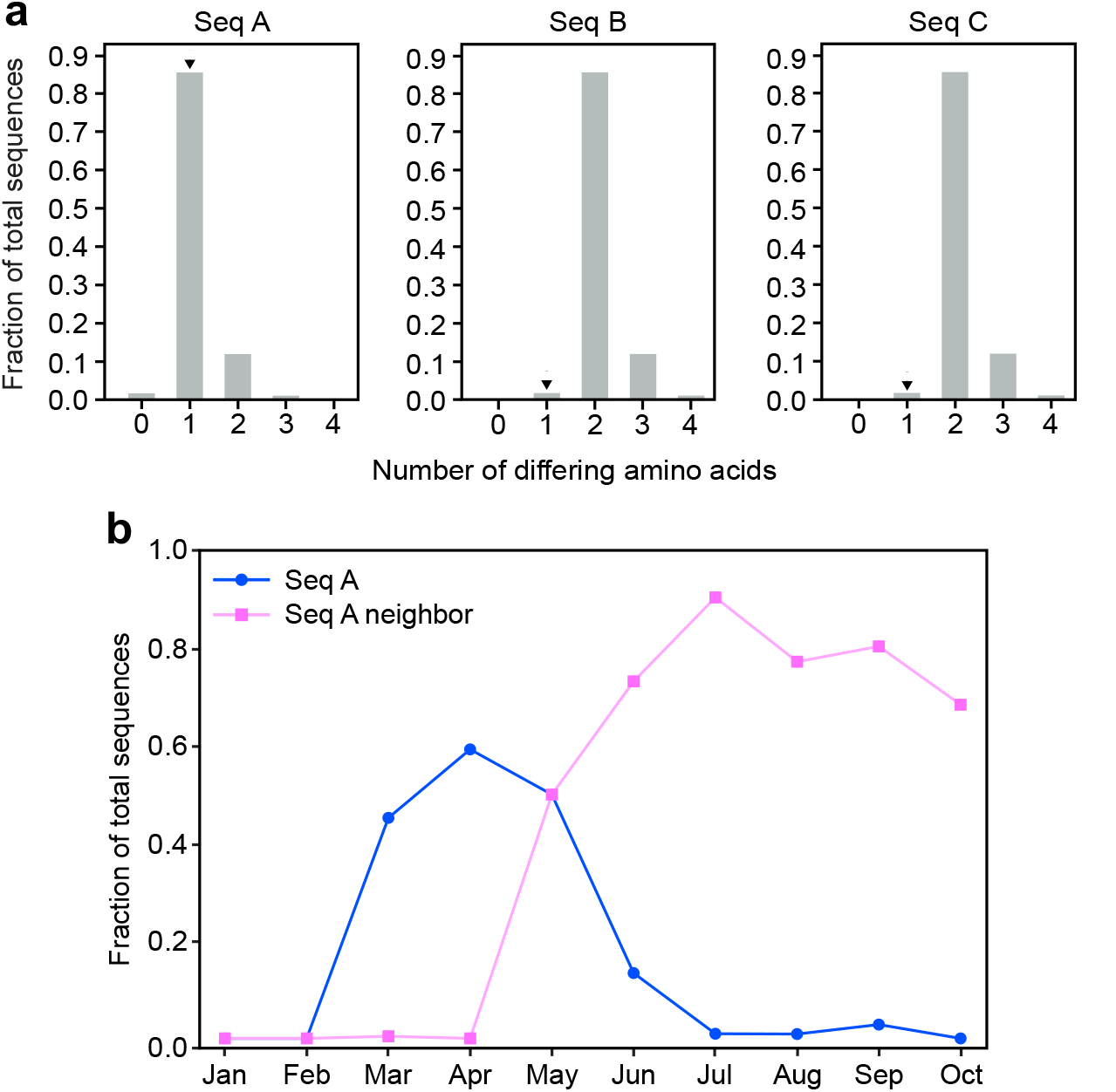
Nearest neighbor analysis of the three most frequent spike protein predictions. (a) Number of amino acid differences between the three most frequent GP predictions and Australian sequences collected within the five-month prediction period (between May 30 and November 30, 2020). Arrowheads indicate the sequence’s nearest neighbors. (b) Monthly frequency of Seq A and its nearest neighbor across the entire Australian spike protein dataset.

## 4. Discussion

### 4.1. Analyzing VPRE’s predictions

VPRE leverages deep learning and GP regression to predict future variants of the SARS-CoV-2 spike protein. We analyzed the first seven months of available SARS-CoV-2 sequences by converting them into continuous latent dimension representations using a VAE, and by performing statistical regression on the continuous data using a GP. Predictions were made by extrapolating the GP model into the future, followed by decoding with the VAE decoder. As a proof-of-concept, we predicted sequences up to five months into the future after our training data cut-off of May 30, 2020, in Australia.

The two most frequent predicted variants, N477S and D614G, were seen in the data that was collected during and one month after the training period, suggesting that no major evolutionary events happened in those 30 days. This finding corresponds with the evolutionary trends we observed in Australia over this period, where no variants rose to dominance within a one-month period. Interestingly, the mutation at position 477 is not a mutational hotspot and may result in lower binding affinity to ACE2 [43], but no data has suggested that this mutation lowers the fitness of the spike protein, so further *in vitro* testing is required to validate the prediction. The other mutation at position 614 is a mutational hotspot that has been previously recorded [44]. In our dataset, D614G is the most prevalent strain. Since our most likely prediction also contained D614G, this suggests that predictions from our model were realistic, at least in recapitulating the evolution of dominant SARS-CoV-2 strains.

Through its continuous numerical approach, VPRE showed potential in modeling viral protein evolution. The VAE was able to translate accurately between numerical coordinates and spike protein sequences, suggesting that it is possible to capture the complexity of amino acid sequences with only three continuous numerical values. The nearest neighbor to our most frequent prediction was only one amino acid different, even though our GP was trained on only 104 amino acid sequences within a timespan of 5 months. Given that our dataset spanned a relatively short timeframe and had limited diversity, it is expected that it was not entirely accurate, especially with the other 16 predictions it generated. Hence, it is reasonable that VPRE predicted variation from the dominant sequences at a very low frequency. It should also be noted that VPRE had no direct measure of antigenic shift or fitness other than the extent of these variables captured in the character of spike protein sequences over time. Therefore, predictions based solely on patterns of amino acid sequence evolution may not be sufficient for a well-informed *in silico* platform. We expect the performance of VPRE to be improved with a more diverse sequence dataset over a longer timeframe.

Further computational and/or experimental validation of VPRE’s predictions would increase the robustness of our model. Because the SARS-CoV-2 spike protein enables host cell entry by interacting with the ACE2 receptor, it would be informative to test the binding affinity of the predicted spike sequences to ACE2 [43]. Quantifying the expression and binding affinity of VPRE’s predicted spike proteins relative to the wildtype SARS-CoV-2 would inform on its fitness, which is one factor that influences the likelihood of the predicted variant emerging and becoming dominant.

### 4.2. Limitations of the model

Although our approach provides a novel way to analyze viral evolution, VPRE’s performance is impacted by limitations in our training dataset. Data-dependent models, especially those under a short time series, are more prone to a high risk of bias [45] and are constrained in their accuracy to the specific protein family they are trained on [26]. Additionally, the dataset was imbalanced across geographical regions and time. Data from North America dominated the training data, which likely biased the VAE’s decoding ability. Most sequences were collected after March 2020, when the virus had already circulated the globe, which posed a challenge in considering the early epidemiology of SARS-CoV-2 evolution. We expect the performance of the model to improve as the scale of SARS-CoV-2 data nears that of endemic viruses such as influenza. Indeed, more coronavirus data is becoming available on a daily basis, and the re-application of existing VAEs to scale to large datasets and predict the effects of mutations on spike protein function would prove useful [22, 46].

We intended to analyze sequences belonging to the same evolutionary clade by first building a phylogenetic tree of spike proteins. However, because the variation between spike protein sequences is limited to a few substitutions or a single amino acid deletion, and most phylogenetic analysis software is optimized for larger genomic variation, we were not able to build an informative phylogenetic tree. To overcome this limitation, we hypothesized that sequences collected from the same geographical region should be subjected to similar evolutionary constraints, and thus be more likely to belong to the same evolutionary path. Hence, the GP model was trained with sequences collected only from Australia. Future attempts to build a phylogenetic tree of spike proteins will likely require whole genome sequencing data.

Working with a small dataset of less than 10,000 sequences which had low levels of variation also posed a challenge for the neural network training. We overcame this issue by synthesizing a dataset of 20,000 spike protein sequences with equally amplified mutation frequencies. This functionally limited the appearance of truly random point mutations in the predictions. While novel sequences and point mutations were predicted, the VAE might have had have a decreased ability to decode point mutations that it had not already observed [47]. Amplifying the mutational variation might have also introduced unnatural features in spike proteins or biased the network’s ability to model variants better than conserved amino acids. Although it is difficult to discern the magnitude of this effect, it appears that the impact was minimal, given that VPRE produced sequences corresponding to real world spike protein evolution in Australia.

The nature of the algorithms we used are such that each variable created by the VAE is fed into an independent Gaussian process. Ideally, all three variables generated by the VAE would be combined in a single multivariate Gaussian interpretation, given their strong interdependence. Interestingly, we found no libraries utilizing integrated tri-variate Gaussian models in open-source repositories or available literature. As such, it seems future work in this stream is needed to construct a more robust multivariate model.

### 4.3. Conclusions

VPRE represents a novel approach to modeling protein evolution using continuous numerical values to encode protein sequences. The software was designed to show the utility of modeling evolution in continuous latent spaces, with the aim of progressing computational sequence prediction and bringing bioinformatics closer to accurate *in silico* protein evolution. Synthetic biology could play a critical role in this process: encoding the predicted spike protein sequences on plasmids would allow them to be readily transferred between model systems and laboratories. For example, a VPRE-predicted spike protein could be expressed in a yeast surface display system to assay ACE2 binding ability. The same predicted spike protein could then be used to pseudotype a non-pathogenic carrier virus and tested for antibody evasion using blood serum samples from vaccinated individuals.

In the process of developing VPRE, we identified areas for continued development, including the need for more powerful phylogenetic reconstruction algorithms and multivariate Gaussian processes. Overall, the implementation of predictive tools such as VPRE opens a window for further investigation into continuous evolutionary models that seek to improve epidemiological modeling, public health intervention systems, and vaccine development.

## Abbreviations

ACE2: angiotensin-converting enzyme 2
COVID-19: coronavirus disease 2019
GP: Gaussian process
SARS-CoV-2: severe acute respiratory syndrome coronavirus 2
VAE: variational autoencoder
VPRE: Viral Predictor for mRNA Evolution

## Acknowledgments

We thank Sibyl Drissler, Avery Noonan, Kristina Gagalova, Carmen Bayly, Arjun Baghela, Alina Kunitskaya, and Evan Gibbard for their feedback and support during the development of VPRE. Katrina Zaraska, Shira Agam, Daniel McClement, Ahmed Abdelmoneim, Kalen Dofher, Katherine Bessai, Mona Golmohammadzadeh, and Morris Huang provided helping hands in the early stages of our research, as part of UBC’s International Genetically Engineered Machine (iGEM) team. This work was funded by the Ecosystem Services, Commercialization Platforms and Entrepreneurship (ECOSCOPE) program and the Professional Activities Fund (PAF) at the University of British Columbia. The funding for ECOSCOPE was provided by the National Sciences and Engineering Research Council of Canada (NSERC) Collaborative Research and Training Experience (CREATE) program.

